# Polygenic adaptation fuels genetic redundancy in *Drosophila*

**DOI:** 10.1101/332122

**Authors:** Neda Barghi, Raymond Tobler, Viola Nolte, Ana Marija Jaksic, Francois Mallard, Kathrin Anna Otte, Marlies Dolezal, Thomas Taus, Robert Kofler, Christian Schlöetterer

## Abstract

The genetic architecture of adaptive traits is of key importance to predict evolutionary responses. Most adaptive traits are polygenic – i.e. result from selection on a large number of genetic loci – but most molecularly characterized traits have a simple genetic basis. This discrepancy is best explained by the difficulty in detecting small allele frequency changes across many contributing loci. To resolve this, we use laboratory natural selection, a framework that is powerful enough to detect signatures for selective sweeps and polygenic adaptation. We exposed 10 replicates of a *Drosophila simulans* population to a new temperature regime and uncovered a polygenic architecture of an adaptive trait with high genetic redundancy among adaptive alleles. We observed convergent phenotypic responses, e.g. fitness, metabolic rate and fat content, and a strong polygenic response (99 selected alleles; mean *s*=0.061). However, each of these selected alleles increased in frequency only in a subset of the evolving replicates. Our results show that natural *D. simulans* populations harbor a vast reservoir of adaptive variation facilitating rapid evolutionary responses. The observed genetic redundancy potentiates this genotypic variation through multiple genetic pathways leading to phenotypic convergence. This key property of adaptive alleles requires the modification of testing strategies in natural populations beyond the search for convergence on the molecular level.

## Introduction

Despite a long-standing interest, the genetic architecture of adaptation is still surprisingly uncharacterized. Traits with simple genetic basis such as pigmentation (*1, 2*), lactose persistence (*3*) and resistance to viruses (*4*), insecticides (*5*) and malaria (*6*) are among the best understood cases of adaptive traits. However, such simple traits are the exception, rather than the rule, and most traits are polygenic, with many contributing loci (*7*-*9*). Selection experiments and QTL studies have identified many loci of small- and large-effects (*10*-*12*). Further support for the polygenic model is provided by GWAS analyses for traits such as human height (*13, 14*), blood lipid levels (*15*), and basal metabolic rate (*16*) which have identified many small-effect loci. However, many causative variants are deleterious typically segregating at low frequencies (*17*). Moreover, for many traits studied by QTL and GWAS the adaptive advantage is not convincingly demonstrated.

Population genetic tests for the identification of selected loci build on predictions for selective sweeps where selection targets independently spread until fixation in the population (*18*). Different allele frequency dynamics are predicted for polygenic traits (*7*). Genomic selection signatures of polygenic adaptation to a new trait optimum have not received as much attention as the sweep model. Most likely because the analysis of extant populations may not be sufficiently powerful. Rather, reliable alternative approaches are needed to identify loci contributing to polygenic adaptive traits. Here, we show that data sets covering multiple time points in replicated populations can distinguish between the allele frequency dynamics of selective sweeps and polygenic traits. We further demonstrate that thermal adaptation is based on an unprecedented high genetic redundancy, with had been predicted (*19*), but has been rarely found (*20, 21*).

## Results

### Increased fitness in the hot-adapted replicates

Temperature is a key environmental factor for all ectotherms and the associated adaptive response in *Drosophila* involves many contributing loci (*22*). To understand the genomic architecture of this canonical polygenic trait, we exposed 10 replicates (each with 1000 individuals) from 202 *Drosophila simulans* isofemale lines to a new hot temperature regime that cycled every 12 hours between 18 and 28 °C, mimicking night and day. After more than 100 generations, we assessed the adaptive response of the 10 evolving replicates. We contrasted fecundity of the ancestral population with each of the evolved replicates after rearing all of them in the hot environment. In agreement with previous results in *D. melanogaster* (*23*), the evolved replicates had significantly higher fecundity, and therefore fitness, than the ancestral population (ANCOVA, *p*<0.0001, Fig. S1A).

### Reconstruction of the selected haplotype blocks

To characterize the genomic signature of adaptation in the evolved replicates, we generated replicated time series data by sequencing pools of individuals (Pool-Seq (*24*)) from the evolving replicates every 10^th^ generation. After stringent filtering steps (Methods), we obtained 5 million SNPs on the major chromosomes. We screened for SNPs with more pronounced allele frequency changes (AFC) following 60 generations of evolution than were expected by genetic drift alone. When combining information across all replicates (Cochran-Mantel-Haenszel, CMH, test) and a single replicate at a time (Fisher’s exact test) we obtained 52,199 candidate SNPs. The number of reported candidate SNPs is likely heavily inflated because these statistical tests assume independence of all SNPs, which is probably violated in our experimental population due to linkage disequilibrium between candidate SNPs (*25, 26*) (in particular for candidate SNPs with a low starting frequency (*27*)). Reasoning that SNPs which are specific to selected haplotypes will have correlated allele frequencies across replicates and time points (*27*), we clustered SNPs by allele frequencies and reconstructed selected haplotype blocks (Fig. 1, SI Text). In total, we identified 99 haplotype blocks, sized between 1.65kb and 5Mb (Fig. S2). We confirmed the accuracy of the reconstructed haplotype blocks by sequencing 100 phased haplotypes from five different evolved replicates and 189 ancestral haplotypes. We compared the SNP markers of the identified haplotype blocks with the phased haplotypes, and 96% of the identified blocks were validated, demonstrating the robustness of our approach (see Fig. 1F for an example, Table S1). For the subsequent analyses each haplotype block is considered a selected allele (SI Text: Definition of selected allele).

**Fig. 1.**
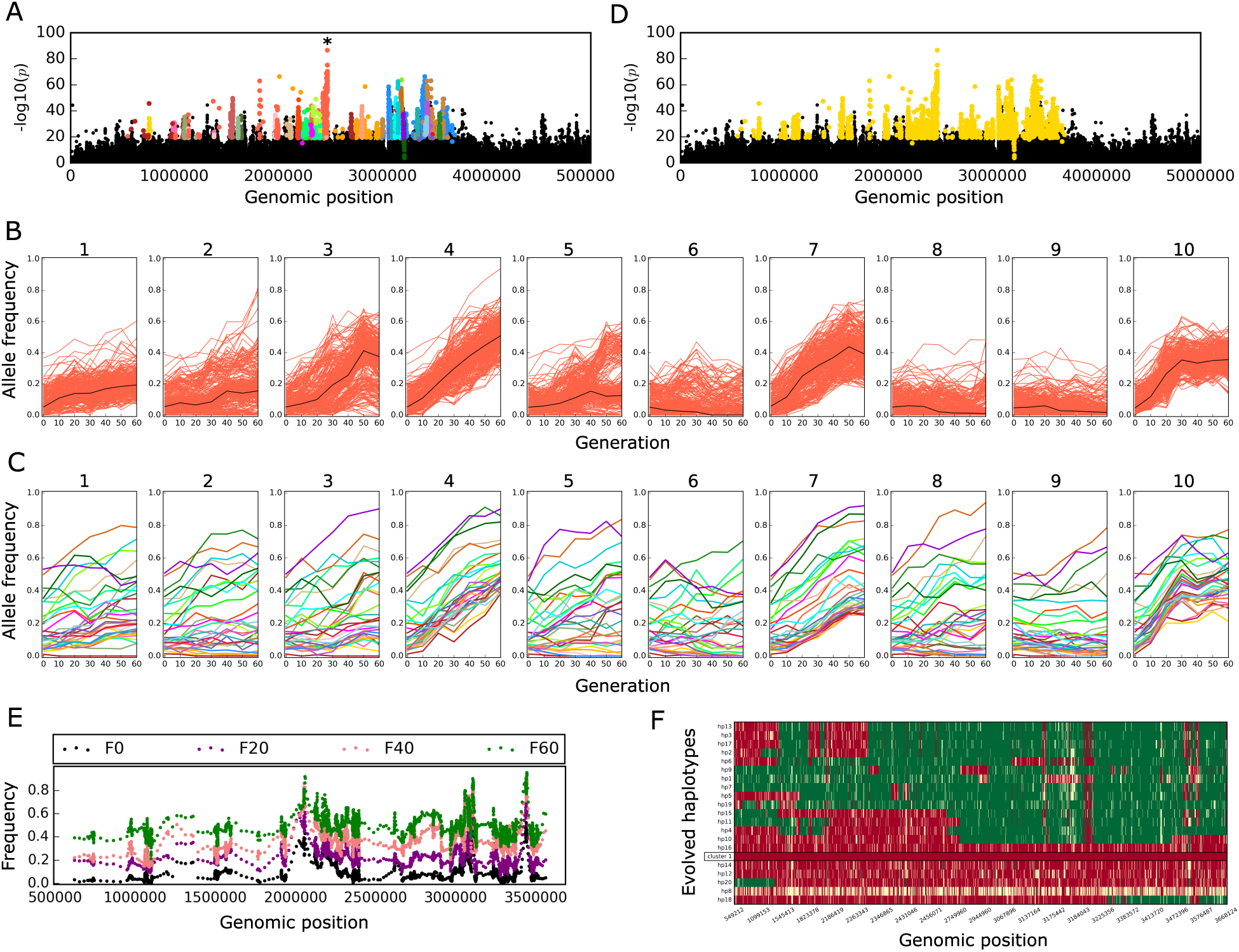
Reconstruction of selected haplotype block from Pool-Seq data. (A, D) show *p*-values from CMH test contrasting the ancestral (F0) with evolved (F60) populations (chromosome 2L: 1-5 Mb). SNPs with highly correlated allele frequencies across replicates and time points are clustered together with stringent clustering (A) and each cluster is indicated by a different color. SNPs in each cluster (e.g. the orange cluster marked with an asterisk in panel A) have correlated frequency trajectories across 60 generations in 10 replicates (B), the black line depicts the median allele frequency trajectory. C) Trajectories of the median allele frequency of the correlated SNPs are shown for each of the clusters in this region (color code corresponds to panel A). Despite different starting frequencies, the median trajectories greatly resemble each other, suggesting that they are correlated and reflect a large selected genomic region. D) Combining the adjacent short clusters with less stringent correlation identifies an allele with weakly correlated SNPs. All SNPs, which cluster together, are used as markers for this selected haplotype block E) Time resolved allele frequencies of marker SNPs for the haplotype block (D) are plotted along their genomic positions. Each dot indicates the mean frequency of 5 SNPs in overlapping windows (offset = 1 SNP). The time-resolved allele frequencies show a consistent increase across the entire haplotype block throughout the experiment. The data are from replicate 4. F) The reconstructed haplotype block (D, E) is experimentally validated. Rows represent 20 phased haplotypes from evolved replicate 4. The reconstructed haplotype block is indicated by a black frame. Each column corresponds to a marker SNP, with red color indicating the character state of the haplotype block, green the alternative allele, and yellow an unknown nucleotide. Missing data are shown in yellow.

### Phenotypic convergence of the hot-evolved replicates

To determine the selected phenotypes underlying the genomic selection signatures, we performed GO enrichment analyses of genes overlapping with the SNP markers of the selected alleles. Consistent with there being many selected genes contributing to similar phenotypes, we detected significant enrichment in several GO categories including oxidative phosphorylation, mitochondrial respiratory chain, ATP synthesis coupled electron transport, melanin biosynthesis process, monosaccharide transportation activity, DNA repair and endopeptidase activity (Table S2). Moreover, the KEGG pathway oxidative phosphorylation was also significantly enriched (Table S3).

The GO enrichment analysis suggests that several phenotypes changed in response to the hot environment. We chose resting metabolic rate and fat content as high-level phenotypes for experimental validation because 1) they both indicate the enrichment for metabolic pathways (oxidative phosphorylation pathway), 2) fat content has been shown to respond to temperature (*28*), and 3) gene expression differences between hot and cold adapted *Drosophila* populations from Africa and Europe also reveal metabolic differences (*29*). Consistent with the genomic signature, both fat content and metabolic rate differed between the ancestral population and the evolved replicates. Females in evolved replicates contained significantly less body fat than the ancestral population (*p*=0.0007, Fig. 2A), and had higher metabolic rates (*p*<0.0001, Fig. 2B), but no difference was noted for males (*p*>0.7, Fig. 2A, B). No significant difference was detected for either of these two high-level phenotypes among the 10 evolved replicates (*p*>0.05 Fig. 2C, D). Thus, the 10 evolved replicates converged not only for fitness (Fig. S1B), but also for other high-level phenotypes, i.e. fat content and resting metabolic rate.

**Fig. 2.**
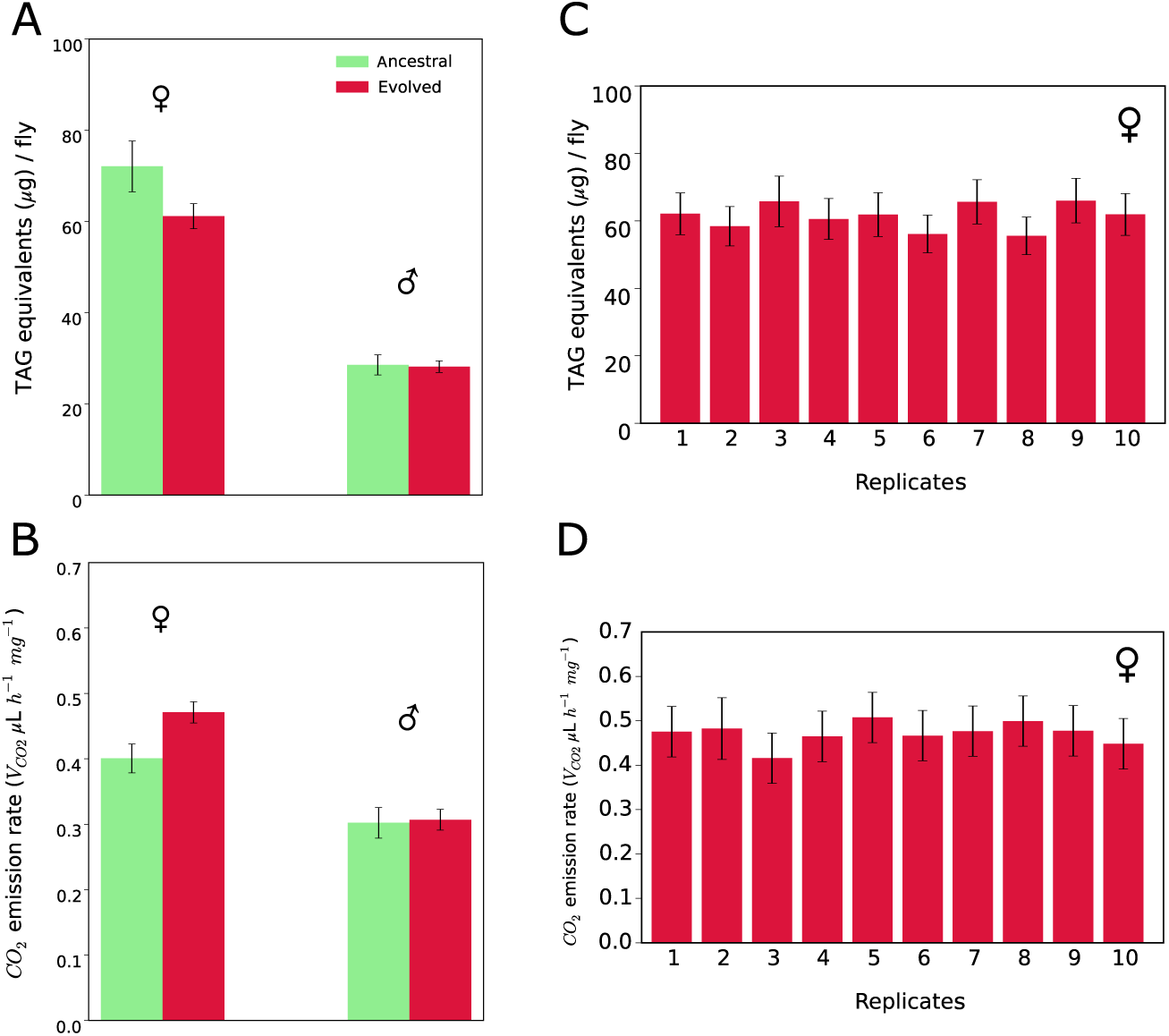
Adaptive response and phenotypic convergence in hot-evolved replicates. A) The amount of triglyceride (TGA) as main constituent of body fat. Females of hot-adapted replicates have significantly lower fat content than the ancestral population. Two-way ANOVA with interaction, Tukey’s HSD test *p=*0.0007, B) Resting metabolism measured by CO^2^ production as measure of resting metabolic rate. Females of hot-evolved replicates produced significantly more CO^2^ than the ancestral population, two-way ANOVA with interaction, Tukey’s HSD test *p*<0.0001. No significant difference was observed among males of the evolved replicates and the ancestral population (body fat: *p*=0.7680, metabolic rate: *p*=0.7405). Convergent evolution for fat content (C) and metabolic rate (D) for females of 10 evolved replicates. No significant difference was detected between the replicates (Two-way ANOVA, Tukey’s HSD test *p*>0.05). The bars show the least-squares means of the linear model and error bars depict 95% confidence levels of least-squares means. Fat content and metabolic rate in males is shown in Fig. S1C and D.

### Genomic heterogeneity among the hot-evolved replicates

Our experimental setting allowed us to empirically quantify several fundamental variables in adaptation genetics, including the starting frequency and distribution of selection coefficients (*s*) for adaptive alleles. Further, the highly convergent phenotypic response across the replicates allowed us to investigate the fundamental question of whether adaptation to a specific stress is driven by the same alleles in all replicates or if multiple alternative genetic routes are possible. Most of the selected alleles started from a low frequency in the ancestral population, but several alleles were rather common with frequencies up to 0.76 (Fig. 3A). The strong selection coefficients of the selected alleles measured across the replicates (min=0.023, max=0.144, Fig. 3B) suggest that a highly parallel genomic architecture could be expected (*30, 31*). However, we observed a highly heterogeneous response across the 10 replicates. A characteristic example is shown in Fig 1B and C, where a selected allele undergoes a striking frequency change in some replicates, but shows no change in others. Indeed, we find that most of the 99 selected alleles increase in only four to six replicates. Few alleles showed a selection response only in one or two replicates and only a single allele increased in all 10 replicates (Fig. 3C-D, Fig. 4B, D). On average, 53 selected alleles were identified per replicate (Fig. 4A).

**Fig. 3.**
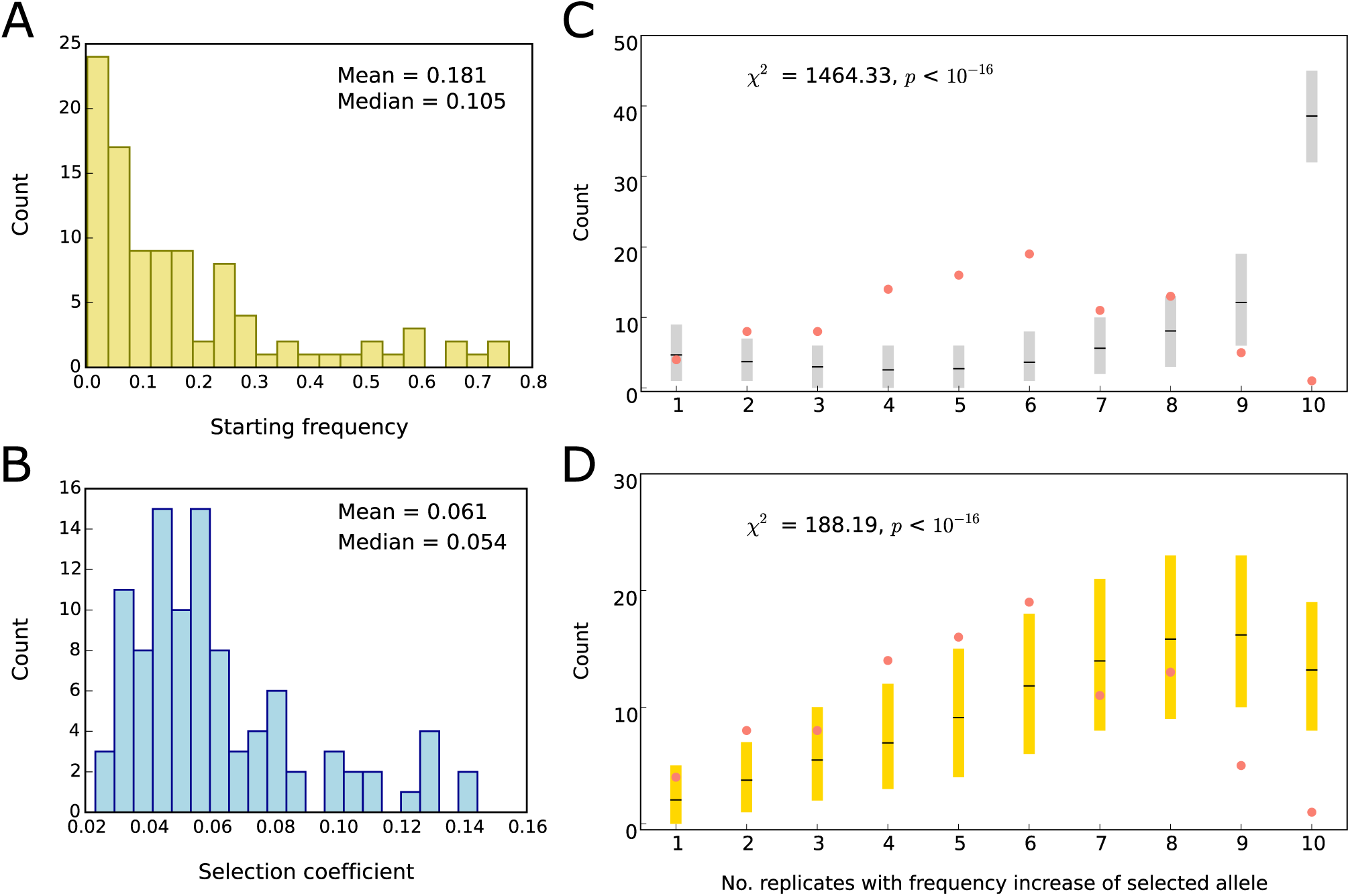
Characteristics of selected alleles: no match to the selective sweep model. Distribution of (A) starting frequencies and (B) selection coefficients (*s*) of the selected alleles. C, D) replicate frequency spectrum, i.e. the frequency distribution of replicates in which selected alleles increase in frequency. An allele with a frequency increase of ≥ 0.1 is considered to be selected in a given replicate. An AFC of 0.1 corresponds to the upper 1% tail of the absolute average AFC distribution across 10 replicates in neutral simulations (SI Methods). The replicate frequency spectrum from experimental data is indicated by salmon dots. The expected distribution of sweep model without (C) and with (D) linkage and constant *s* across replicates using empirical *N*_*e*_ and locus-specific *s* (B) and starting frequency (A) was obtained from computer simulations (Methods). Grey and gold bars show the 95% confidence interval (CI) of 1000 iterations. Means are depicted in black lines in each bar. The χ^2^ goodness of fit tests indicate that selective sweep models do not fit the replicate frequency spectrum from the empirical data.

**Fig. 4.**
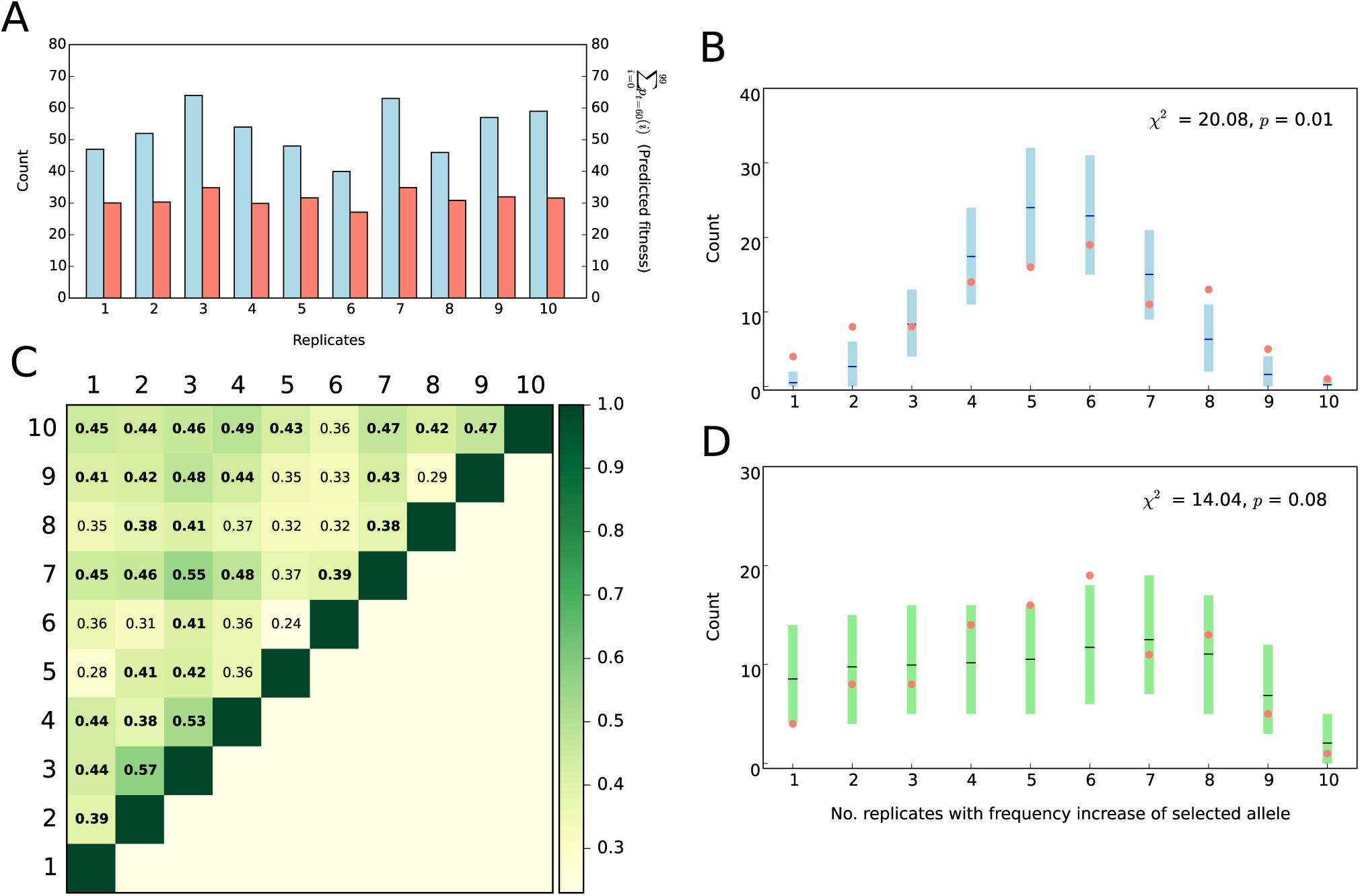
Adaptive genetic redundancy of selected alleles. A) Despite the different number of selected alleles with ≥0.1 AFC (blue bars, average 53 loci/replicate) after 60 generations in each replicate, the sum of frequencies of all 99 alleles (which approximates fitness) at generation 60 is similar (χ^2^ =1.519, *p*=0.99) across replicate (salmon bars). B) comparison of the empirical replicate frequency spectrum (salmon) to the expected one under the assumption of genetic redundancy among alleles. 95% CI (blue) and means (black) were obtained by 1000 iterations of delete-*d* jackknifing (Methods). C) Pairwise Jaccard similarity indices among 10 evolved replicates. Indices significantly higher than expected by chance (Fig. S6) are shown in bold. D) comparison of the empirical replicate frequency spectrum (salmon) to the expected one assuming a quantitative trait model with stabilizing selection after a change in trait optimum. 95% CI (green) were obtained by 1000 iterations of forward simulations (Methods). The χ^2^ goodness of fit tests indicate a much better fit of the genetic redundancy and quantitative trait models.

### Genomic heterogeneity does not match the sweep paradigm

We used these patterns to discern several different adaptive scenarios (Fig. S3). Since the P-element is spreading in the ancestral population in our experiment (*32*) the observed heterogeneity among replicates might have been driven by the new replicate-specific P-element insertions. Nevertheless, a careful examination of P-element insertions showed that the observed insertions occurred at too low frequencies to explain the adaptive response (Table S4).

With most selected alleles starting from a low frequency in the ancestral population (Fig. 3A) and the moderate *N*_e_ of the experimental populations (<300), combination of selection and genetic drift could have contributed to the observed heterogeneity through differential loss of rare selected alleles across replicates. To test if the combined effect of the sweep paradigm and drift could produce the observed heterogeneity among replicates, we simulated evolution under a standard sweep model with constant *s* (different among alleles but similar across replicates and time), but without linkage and epistasis. Using replicate-specific *N*_e_ estimates, and allele-specific *s* and starting frequencies, we simulated 1000 iterations of 99 independent alleles in 10 replicates across 60 generations. We detected some heterogeneity among replicates, but unlike the experimental data most alleles spread in 9 or 10 replicates (Fig. 3C). Importantly, the poor fit to the sweep model could not be explained by how *s* was estimated, as the same trends are seen regardless of which estimation procedure was used (Fig. S4, S5).

To rule out the possibility that Hill-Robertson interference caused the observed heterogeneity, we included linkage in the standard sweep model using 189 phased ancestral haplotypes. We simulated 99 selected alleles in 10 replicates of a population of 300 diploids (estimated *N*_*e*_) in 1000 iterations with allele-specific *s* and starting frequencies and the recombination rate estimated from the ancestral haplotypes. Despite improving the fit to the experimental data (Fig. 3D), the differences are still substantial and we conclude that the standard population genetic model of selective sweep, including Hill-Robertson interference, cannot explain the heterogeneous distribution of selected alleles among our replicates.

### Adaptive genetic redundancy of the selected alleles

Other factors, such as linkage between selected and deleterious alleles, or frequency-dependent selection, could contribute to the observed replicate heterogeneity. Nevertheless, the combination of the striking phenotypic convergence (Fig. 2C,D) and different subsets of selected alleles across replicates (Fig. 4A) strongly suggests that the ancestral population contained more beneficial alleles than were needed to achieve optimum fitness (*19, 33, 34*), i.e. genetic redundancy. The simplest form of genetic redundancy is when all beneficial alleles have equal effects but not all of them are required to reach the fitness optimum; different combinations of alleles can reach the same fitness in different replicates and subsequently produce a heterogeneous genomic pattern among them. Assuming equal effects of the selected alleles, we attempted to predict the fitness based on the summed frequencies of the selected alleles at generation 60. Similar to fecundity, fat content and metabolic rate, all replicates also converged for our predicted fitness (Fig. 4A), supporting the assumption of genetic redundancy and similar effect sizes of the selected alleles. We further scrutinized genetic redundancy through jackknifing, and randomly sampled a subset of the 99 selected alleles that matched the observed number of selected alleles with ≥0.1 AFC for each replicate (Fig. 4A). This simple model of genetic redundancy fits the observed pattern of heterogeneity among replicates quite well (Fig. 4B). Nevertheless, the combination of selected alleles shared across the replicates was significantly more similar in the experimental data (Fig. 4C, median Jaccard index=0.41) than for randomly combined alleles of the redundancy model (median Jaccard index=0.36, p<0.001, Fig. S6). Despite a significant difference, the experimental Jaccard index is only slightly higher than for random combinations of alleles. Higher similarity of replicates in experimental evolution could be simply because alleles with higher starting frequency increase in frequency in more replicates (Fig. S7A). Thus, unlike a previous report of convergent adaptation in two tree species (*20*), we find no evidence for strong genetic constraints that limit the possible combination of beneficial alleles in our experimental populations. Because our redundancy test did not model the frequency trajectory of selected alleles, we also simulated a simple quantitative trait model with stabilizing selection. Assuming the same effect size for all 99 alleles in 10 replicates with starting frequencies matching the experimental data, this simple quantitative trait model nicely matched the observed heterogeneity pattern (Fig. 4D).

## Discussion

Quantitative genetic theory assumes that in populations close to the trait optimum, strong effect alleles do not segregate at intermediate frequencies (*35, 36*). Thus, after shifts of the trait optimum, the phenotypic response is typically mediated by many small effect alleles with no discernable change in allele frequency (*7*). In our data, strongly selected alleles (mean *s*=0.061) do occur at intermediate frequencies (Fig. 3A, max=0.76, mean=0.18). This discrepancy may have several explanations, ranging from an ancestral population that has not reached the trait optimum to major impact of genetic drift and population structure. The abundance of large effect alleles contributing to adaptation also contradicts another theoretical prediction that polygenic trait adaptation is driven by alleles of small effect (*37*).

With 99 selected alleles and 202 founder isofemale lines, on average every second founder carries a different selected allele. This implies that natural *D. simulans* populations harbor vast reservoirs of variants capable of contributing to temperature adaptation and different combinations of these variants result in similar phenotypic changes. Thus, beneficial alleles tend to segregate at higher frequencies than neutral SNPs in our experimental population (Fig. 5), suggesting a role for balancing selection – possibly driven by seasonal temperature changes.

**Fig. 5.**
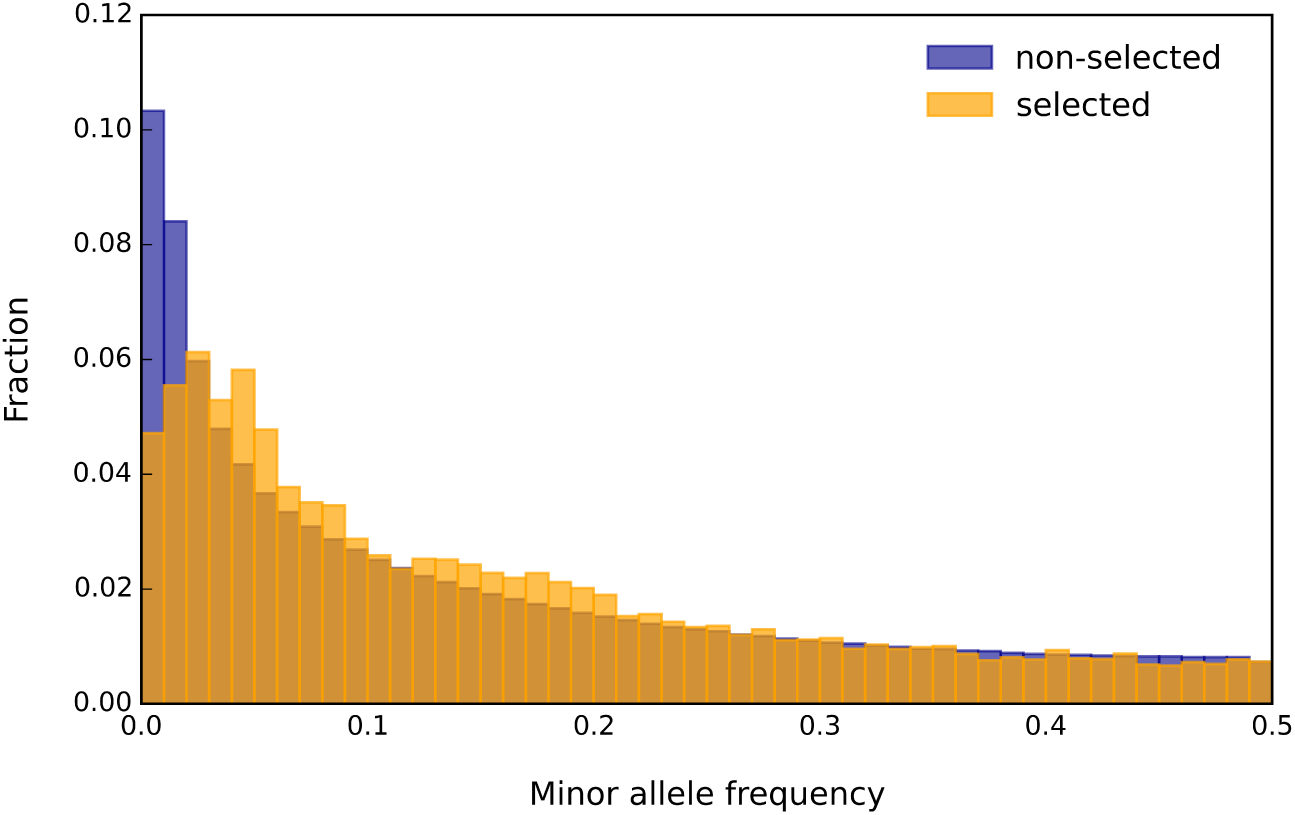
Selected alleles occur at higher frequency than neutral SNPs. The folded site frequency spectrum (SFS) of marker SNPs from selected alleles (23,835 SNPs) and non-selected SNPs (5,072,365 SNPs) is plotted. SNPs that were not identified as markers of selected alleles are considered non-selected SNPs. The distributions of these two SNPs classes differ significantly (two sided Kolmogorov-Smirnov test, ks value = 0.086, *p* < 10^−16^).

Is the adaptive genetic redundancy of temperature adaptation an exception or do more traits have a similar genetic basis? The empirical evidence for adaptive genetic redundancy is extremely sparse, but this probably reflects a bias toward methodologies that search for convergent genetic changes. Examples for adaptive genetic redundancy are *de novo* mutations of experimental *E. coli* populations (*38*), pigmentation in African *Drosophila* where different genes contribute to the same dark phenotype (*39*), desiccation resistance (*21*), and the hemoglobin oxygen affinity mediated by different amino acid substitutions in 56 avian taxa (*40*). Truncating selection studies in *Drosophila* (*41*) and corn (*42*) reporting rapid phenotypic responses despite a small number of founders, also indirectly support the presence of abundant genetic redundancy in natural populations. The abundance of large effect beneficial alleles segregating in natural populations ultimately means that there are many genetic pathways to the same fitness optimum. Thus, the genomic signature of adaptation could differ among natural populations, and genome scans for convergent genomic signatures across populations are less likely to succeed for such quantitative traits.

## Methods

### Experimental setup and reconstruction of the selected haplotype blocks (selected alleles)

Ten replicate populations were set up using 202 isofemale lines from a natural *D. simulans* population collected in Tallahassee, Florida, USA (*43*). The replicates, each with a population size of 1000, were maintained in a new hot environment alternating between 18 and 28°C every 12 hours. All replicates were subjected to Pool-Sequencing at generation 0 and every 10^th^ generation until generation 60. After stringent filtering steps (SI Methods), we identified SNPs, which changed more in allele frequency than expected by neutral drift. To account for LD we shift the focus from SNPs to selected haplotype blocks which carry the selected target(s) using a modification of a recently published approach (*27*). A detailed explanation of candidate SNP inference and haplotype blocks reconstruction is provided in SI Methods. Details on the pipeline for identification of transposable elements and Gene Ontology (GO) and pathway enrichment analyses are provided in SI Methods.

### Simulations for selective sweep and quantitative genetics model

We used computer simulations to discern several different adaptive scenarios (Fig. S3). More details on these simulations are provided in SI Methods.

#### A. Sweep model without linkage and a constant selection coefficient across replicates

Forward Wright-Fisher simulations were performed using 99 independent alleles (matching the observed number of selected alleles), the observed starting frequency (median frequency of SNPs characteristic to the selected haplotype block/allele) in the ancestral replicates (Fig. 3A), replicate-specific *N*_*e*_ (Table S5) and allele-specific *s* (Fig. 3B), assuming no linkage and epistasis. Simulations with different parameters were performed to assure that the results are not biased by the allele-specific starting frequency and *s* (Fig. S5).

#### B. Sweep model with linkage and a constant selection coefficient across replicates

We used 189 individually sequenced haplotypes from the ancestral population (SI Methods) for the simulations. We simulated 99 selected alleles in 10 replicates of a population of 300 diploids (corresponding to the estimated *N*_*e*_) with the recombination rate estimated from the haplotypes. The allele-specific *s* (Fig. 3B) and starting frequency (Fig. 3A) were used for simulations.

#### C. Genetic redundancy model

Assuming that all alleles are functionally equivalent, we generated 1000 data sets using delete-*d* jackknifing. In each set the number of selected alleles for each replicate matched our observations (Fig. 4A), but was randomly drawn (without replacement) from the total of 99 selected loci.

#### D. Quantitative trait model

Using forward simulations, we simulated a quantitative trait after a change in trait optimum in 10 replicates with 99 contributing alleles using the empirical starting frequency (Fig. 3A). Alleles were in linkage equilibrium and had equal effects. A good fit to the empirical data was obtained with fitness ranging between 0.5-4.5 and a mean fitness optimum of 0.6±0.3 (standard deviation).

### Phenotypic assays

**Fecundity:** The flies used for phenotypic assays were maintained at controlled density in a common garden for at least 2 generations. The fecundity of evolved and ancestral populations each with three to six technical replicates was estimated by counting the total number of eggs laid by females during four days (between day two to five after eclosion). **Resting metabolism assay:** Females and males were four to five and six to seven days old, respectively, during resting metabolic measurements. For each sample the resting metabolic rate of 150 flies was measured by repeated CO^2^ emission measurements of stop-flow respirometry (Sable Systems, Las Vegas, Nevada, USA). **Body fat assay**: The body fat was measured in four days old females and males. Homogenates were prepared as described in (*44*) and lipid measurements were performed by coupled colorimetric assay as described in (*45*). To test for the significance between the ancestral and evolved (all 10 evolved replicates combined) populations and also between the 10 evolved replicates, the results of phenotypic assays were analyzed using linear model. More details on the phenotypic assays and statistical tests are provided in SI Methods.

## Data availability

The raw reads for Pool-Seq libraries of all replicates and the raw reads for haplotypes are available from the European Sequence Read Archive under the Accession numbers SRAxxxxxxxxxxx. SNP data sets and scripts are available from the Dryad Digital Repository under xxxxxxxxxxxxxx.

## Acknowledgements

We thank Nick Barton, Reinhard Bürger, Joachim Hermisson and Ilse Höllinger for comments on an earlier version of manuscript. We thank Franziska Hofstetter for helping with the fat content assay. We are grateful to members of the Institut für Populationsgenetik for helpful discussions. This work was supported by the European Research Council grant “ArchAdapt” and the Austrian Science Fund (FWF, W1225-B20). K.A.O. was supported by a DFG Research Fellowship (OT 532/1-1) and F.M. was supported by a Marie Skłodowska Curie Individual Fellowship (H2020-MSCA-IF-661149). T.T. is recipient of the DOC fellowship of the Austrian Academy of Sciences.

## Author contributions

C.S. and N.B. designed the study. R.T. collected *D. simulans* flies and established the experimental evolution populations. V.N. generated and managed the NGS data. N.B. analyzed the data. N.B., A.M.J., K.A.O. and F.M. performed fitness assay. N.B. performed fat content and metabolic rate assays. M.D. supervised the statistical analysis. R.K. provided the simulation tool. T.T. provided ancestral haplotypes and *D. simulans* recombination rates. N.B. and C.S. wrote the manuscript with input from coauthors. All authors approved the final manuscript.

